# Early Diabetic Ca^2+^ Handling Impairments in the Rod Bipolar Pathway

**DOI:** 10.64898/2025.12.01.691679

**Authors:** Jared T. Hill, Andrea J. Wellington, Diego S. Del Villar, Erika D. Eggers

## Abstract

**Background:** Previous work showed that electrically-evoked inhibition to Rod Bipolar Cells (RBC) is reduced in a mouse model of early diabetes. It is hypothesized that this is due to impaired Ca^2+^ handling in the presynaptic amacrine cell, either through increased Ca^2+^ buffering or decreased influx. To test this hypothesis and develop a mechanism for this effect, a model where direct optogenetic activation of inhibitory amacrine cells that expressed the light-activated channel ChR2 was used to isolate amacrine cell inputs to RBCs. Application of selective Ca^2+^ channel blockers could then assess potential locations of amacrine Ca^2+^ disruption. Using whole cell patch clamp electrophysiology, recordings were made from a 6 week diabetic population (DM) and vehicle injected non-DM animals.

**Results:** Robust GABA_C_ receptor inhibitory currents were recorded from RBCs after ChR2 stimulus that were significantly diminished by the application of nifedipine to block L-type Ca^2+^ channels in both DM and non-DM conditions. There were significant differences in the peak amplitude of these responses between DM and non-DM groups (p = 0.0146). However, in the non-DM group the decay tau of the response to the 50ms stimulus was significantly diminished by nifedipine (***τ*** p =0.0498, n = 5), but this was not seen in the DM group (***τ*** p = 0.9498, n=7). A 1s nifedipine-reduced response saw its decay tau increase in the DM group but not the non-DM. Ca^2+^ - induced Ca^2+^ release (CICR) blockade with ryanodine decreased responsivity equally between groups in the 1s stimulus but showed no significant kinetic changes. CICR blockade for a 50ms stimulus response showed significant kinetic changes in diabetes but otherwise reduced the response equally between DM and non-DM. Blockade of the mitochondrial Ca^2+^ uniporter (MCU) had little effect on the optogenetic response.

**Conclusion:** This study presents evidence that diabetes alters amacrine cell output to the RBC unmasked through blockade of the L-type calcium channel, and the Endoplasmic Reticulum (ER). An apparent explanation for our results is that DM calcium buffering is dysregulated, leading to prolonged responses. The underlying mechanism for this alteration is complex and not yet clearly elucidated.

## Introduction

Diabetic retinopathy (DR) is a common complication of diabetes and the leading cause of blindness in the working age population^1–4^. Evidence is mounting that early DR produces neural dysfunction before the classical overt vascular pathology, with particularly consistent changes in dim-light (rod-pathway) signaling^5–7^. Human and animal studies show altered electroretinographic timing and amplitude in diabetes (DM), especially in oscillatory potentials (OPs)^7^. OPs largely reflect amacrine–bipolar cell interactions, with subsequent alteration indicating inner-retinal circuit disruption early in disease^8–10^. These functional abnormalities include delayed OP implicit times suggesting altered or reduced inhibitory drive within the ON pathway^11^. Excessive activity of the ON pathway was reported where 3-4 months of STZ induced diabetes increased ON ganglion cell spiking rates in DM animals^12^, due to impaired inhibition in the ON pathway. Light-evoked inhibition to rod bipolar cells is also decreased after 6 weeks of diabetes^13^. The mechanisms of this inhibitory disruption are unclear, yet research is progressing towards identifying specific circuit level contributors to this dysfunction.

In the healthy retina, GABAergic and glycinergic amacrine cells provide inhibition to Rod Bipolar Cells (RBC) through three primary receptors: glycine receptors (GlycineRs), fast GABA_A_ receptors (GABA_A_Rs), and slow decay GABA_C_Rs. Isolation of these distinct pathways showed that GABA_C_Rs mediate the largest component of light evoked inhibitory charge transfer, while GABA_A_Rs and GlycineRs make smaller, fast kinetic contributions^14^. The inhibitory amacrine synapses onto the RBC release neurotransmitter for hundreds of milliseconds past the offset of a stimulus, in asynchronous release with slow presynaptic Ca^2+^ dynamics^14–16^. L-type Ca^2+^ channels provide the bulk of the Ca^2+^ entrance, with the signal also prolonged by CICR^15, 17^. We previously showed that after 6 weeks of diabetes, these Ca^2+^ dependent release characteristics are disrupted^16^. DM GABA_C_R isolated electronically activated inhibitory post-synaptic currents (eIPSCs) were significantly more diminished when exposed to the Ca^2+^ buffer EGTA-AM than their non-DM counterparts. This suggests that diabetes is reducing either the capacity of Ca^2+^ to enter the cell, its ability to be released from internal stores, or that Ca^2+^ is more strongly buffered in diabetes. This present work builds on these studies by using optogenetics to precisely activate all RBC inhibitory synaptic partners and thereby isolate these inputs. By pairing this optogenetically induced IPSC (oIPSC) with selective Ca^2+^ channel blockade we aim to determine which components of Ca^2+^ handling are altered in diabetes

## Methods

### Animal model

B6.Cg-Tg(Slc32al-COP4*H134R/EYFP) male mice^18^, which express ChR2 under the VGAT promoter in the inhibitory amacrine and horizontal cells, were used for all experiments according to protocols approved by the University of Arizona Institutional Animal Care and Use Committee. Animals were housed in the University of Arizona animal facility and given the National Institutes of Health-31 rodent diet food and water ad libitum.

### Induction of Diabetes

7 week old mice received 3 consecutive, once daily, 75mg/kg streptozotocin or citrate vehicle injections intraperitoneally after fasting for 4 hours. Animals were monitored for health and weight loss, and their urine was tested weekly to confirm glycosuria, classified as greater than 2000 mg of glucose / dl. After 6 weeks of diabetes the mice were fasted for 4 hours, and blood tested via the OneTouch UltraMini blood glucose monitoring system (OneTouch UltraMini, LifeScan; Milpitas, CA, USA). Diabetes was assessed as having a blood glucose reading over 250 mg/dl.

### Solutions and drugs

A modified ringer’s solution was used for the extracellular bath and as the dissection solution, bubbled with 95%/5% O₂/CO₂, and contained (in mM): 125 NaCl, 2.5 KCl, 1 MgCl₂, 1.25 NaH₂PO₄, 20 Glucose, 26 NaHCO₃, 2 Na-Pyruvate, 4 Na L-Lactate, .5 L-Glutamine, 2 CaCl₂. The intracellular solution used for measuring inhibitory currents contained (in mM): 120 CsOH, 120 Gluconic Acid, 1 MgCl₂, 10 HEPES, 10 EGTA, 10 TEA-Cl, 10 Phosphocreatine-Na₂, 4 Mg-ATP, .5 Na-GTP and 50 µM Alexa Fluor 488 (Invitrogen, Carlsbad, CA, USA) with pH adjusted to 7.2 by CsOH. Osmolarity was measured at 320. Unless otherwise noted, all drugs were acquired from Sigma Aldrich. Photoreceptor inputs were pharmacologically blocked with CNQX (25 µM), APV (50 µM), ACET (1 μM), and L-AP4 (50 µM). GABA_C_Rs were isolated by blocking GlycineRs (Strychnine 1 µM) and GABA_A_Rs (SR-95531 20µM). For pharmacological experiments, L-type Ca^2+^ channels were blocked by Nifedipine (10µM). RyanodineRs were blocked by Ryanodine (50µM) (Santa Cruz biosciences). Mitochondrial Ca^2+^ Uniporter (MCU) was blocked by MCU-i4(10µM) (Tocris biosciences).

### Electrophysiology recordings and analysis

After 6 weeks of diabetes, mice were euthanized with CO_2_ and then their eyes were enucleated and placed in a 95% O_2_ / 5% CO_2_ carbogen bubbled Ringer’s solution for dissection. Retinas were dissected out, trimmed, mounted on nitrocellulose paper, cut into 250µm thick slices, turned 90° to have access to all retinal layers, then placed in a carbogen infused dark-box. Whole-cell voltage clamp recordings were made from RBCs held at 0 mV, the reversal potential for non-selective cation channels. For all recordings, borosilicate glass electrodes (World Precision Instruments, Sarasota, FL, USA) had resistances of 7 to 11 MΩ and the series resistance during recordings was typically 10 to 20 MΩ. Liquid junction potentials of 20 mV were corrected prior to recording. Responses were filtered at 5 kHz using a four-pole low-pass Bessel filter on an Axopatch 200 B amplifier (Molecular Devices, Sunnyvale, CA, USA). The response was digitized at 10 kHz using a Digidata 1440A data acquisition system (Molecular Devices) and Clampex software (Molecular Devices). ChR2-expressing cells were stimulated with a 1s and 50ms full-field light stimulus projected through a 60x objective (λ = 470 nm, ThorLabs LED) (Moore-Dotson 2019).

For all cell recordings a 1s stimulus was used (long stimulus), and for a subset a 50ms stimulus was added (brief stimulus) with 20 second rests between stimuli. 1s maximally activated the cell and produced a plateau current. The 50ms stimulus was chosen as it offered a strong and consistent response (superior to the 10ms) but did not lead the current to plateau (100ms +), while still recapitulating the time course of natural physiological responses which can be relatively prolonged^14^.

Each recording used at least two stimulus sweeps which were averaged together for the final analysis. First a cellular baseline recording was performed just in the presence of Ringer’s modified solution. The photoblocking and GABA_C_R isolation solution, containing CNQX, ACET, APV, LAP4 for photoreceptor glutamate blockade, and strychnine + SR-95531 for GABA_C_R isolation, was washed on for 5 minutes. Following the wash on of the GABA_C_R isolation solution, oIPSC responses were recorded. Finally, the experimental condition drug was added to the bath and washed on for 5 minutes, after which the final recording was taken. RBC morphology was confirmed after recording using Alexa fluorescence for cells with somas in the inner nuclear layer and a characteristically long straight axon that ends with large lobules near the ganglion cell layer.

Traces were filtered using a 500 Hz Bessel filter, baselined, then sweeps were averaged together to create a waveform for the baseline condition, the GABA_C_R isolation condition, and the experimental condition. The charge transfer (Q), peak amplitude (PA) and Decay Tau (***τ***) were analyzed for all evoked responses and paired t tests were performed. 50ms responses had their Time to Peak (TtP) measured from onset of stimuli. Data was normalized to the GABA_C_R isolation, and t-tests were performed. Decay Tau was measured from the offset of stimulus in the 1s condition and from the peak of the response in the 50ms condition. Welch’s t tests were performed on any comparisons between DM and non-DM groupings. Baseline and GABA_C_R data were aggregated across experiments for baseline comparisons between non-DM and DM conditions.

### Immunohistochemistry, Imaging and Analysis

Immuno Histochemistry (IHC) was performed with stains for DAPI, GAD65/67, and CAV1.3 to stain L-type Voltage Gated Calcium channels. After 6 weeks post injection, mouse eyes were enucleated, and eye cups (with cornea & lens removed) were fixed for 30 minutes in 4% paraformaldehyde. Eye cups were rinsed in Phosphate Buffered Saline (PBS), then retinas were dissected out. Retinas were embedded in 5% agarose and sliced at 60 µm on a vibratome. Retina slices were transferred to a 24-well dish and incubated for 1 hour in blocking solution (5% goat serum, 10% Bovine Serum Albumin, 0.5% Triton X-100 in Phosphate Buffered Saline). Retina slices were incubated overnight at room temperature in blocking solution containing the following primary antibodies: Mouse IgG1 Anti-CaV1.3 L-type Ca channel 1:200 (DSHB N38/8), Mouse IgG2a Anti-GAD65 1:1000 (DSHB GAD-6) and Mouse IgG2a Anti-GAD 67 1:1000 (Millipore MAB5406). Slices were washed 3×30 min in PBS, then incubated with the following secondary antibodies for 2.5 hours: Anti-mouse IgG1-AlexaFluor555 1:1000 (Invitrogen A21127) to detect CaV1.3 and Anti-mouse IgG2a-AlexaFluor488 1:1000 (Invitrogen A21131) to detect GAD65 & 67. Nuclei were stained with DAPI (4′,6-Diamidine-2′-phenylindole dihydrochloride) 1:1000 (Invitrogen #D1306) during secondary antibody incubation. Slices were washed 3 x 30 minutes in PBS, then mounted in ProLong Glass (Invitrogen #P369820). Slides were imaged on a Zeiss LSM 880 inverted confocal microscope using a 40x objective (Plan-Apochromat 40x/1.3) with 2 x zoom. Z-stacks were acquired using 0.38um slice intervals. Images were collected as 16-bit, 212.55 x 212.55 microns (1024 x1024 pixels). The following laser intensities were used: GAD6567 488nm laser at 1.8%, CaV1.3 561nm laser at 3%, and DAPI 405nm laser at 0.5%.

ImageJ/Fiji was used for image analysis. Max intensity Z-projections were made from 5 sequential optical slices from each vibratome slice. Two to four vibratome slices were used from each retina, except for control retina #684, which had only one usable vibratome slice. Measurements were averaged for each retina. Z-projections were background-subtracted (rolling ball, 10px) and median filtered (3px). A region of interest (ROI) was drawn around each Inner Plexiform Layer (IPL), and the area outside of this was cleared. Thresholds were set at 11.28% of maximum intensity for CaV1.3, and at 16.79% of maximum intensity for GAD 65 &67. The threshold values used showed minimal signal in slices processed without primary antibodies. Thresholded images were measured, then multiplied against each other to find overlapping areas. Student’s t-test with equal variance was used to compare groups.

## Results

### Long stimuli GABA_C_R Isolated Responses are Not Differentially Affected by DM

To test the hypothesis that the source of amacrine cell Ca^2+^ reduction in early diabetes relates to alteration of the L-type, long duration, voltage gated Ca^2+^ channels, recordings were made from patch clamped RBCs with optogenetically invoked inhibition (oIPSC) from presynaptic amacrine cells being measured. Our results show a robust and prolonged response to the 1s stimuli, with variable time to peaks, and an immediate initiation of decay following stimuli offset (Figure 1A-B, Table 1). In the baseline response, small, sharp, glycinergic and/or GABA_A_R currents ride atop the large GABA_C_R mediated current. With application of strychnine and SR 95531 (glycine and GABA_A_R antagonists), these sharp currents disappear, and the total charge transfer becomes significantly larger in non-DM and in DM groups (Figure 1C). Interestingly, the peak amplitude is not significantly affected by GABA_C_R isolation in either experimental group (Figure 1D). Given that GABA_C_R isolation blocks a significant component of serial inhibition, these larger responses were expected.

**Figure 1.**
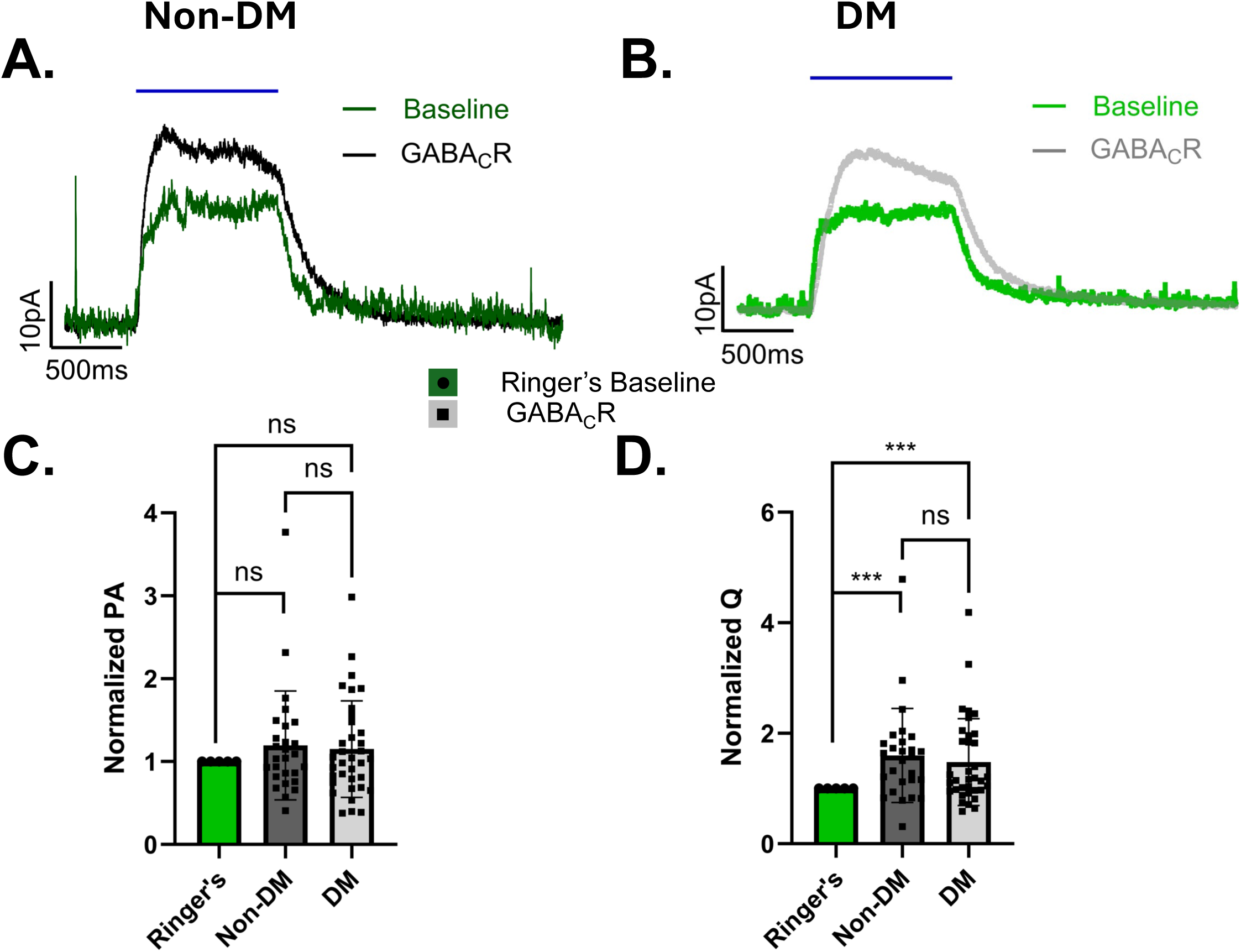
A, Averaged non-DM trace showing baseline Ringer’s recording in dark green, and GABA_C_R isolation in dark grey. B, Averaged DM trace showing baseline Ringer’s recording in light green, and GABA_C_R isolation in light grey. The blue bar represents the 1 second blue light stimulus for both. C, Peak Amplitude in the non-DM group trends larger, but is not significant (PA + 19.6% +/-12.7%, p = 0.1331, n = 27) and shares this trend in the DM group (PA+15.1% +/- 9.8%, p = 0.1337, n = 35). D, both non-DM (Q + 59.6%, p = 0.0012, n = 27) and DM (Q + 47.5%, p = 0.001, n = 35) groups have significant increases in charge transfer following GABA_c_R isolation. All values are normalized to the value in normal Ringer’s solution.

**Table 1:**
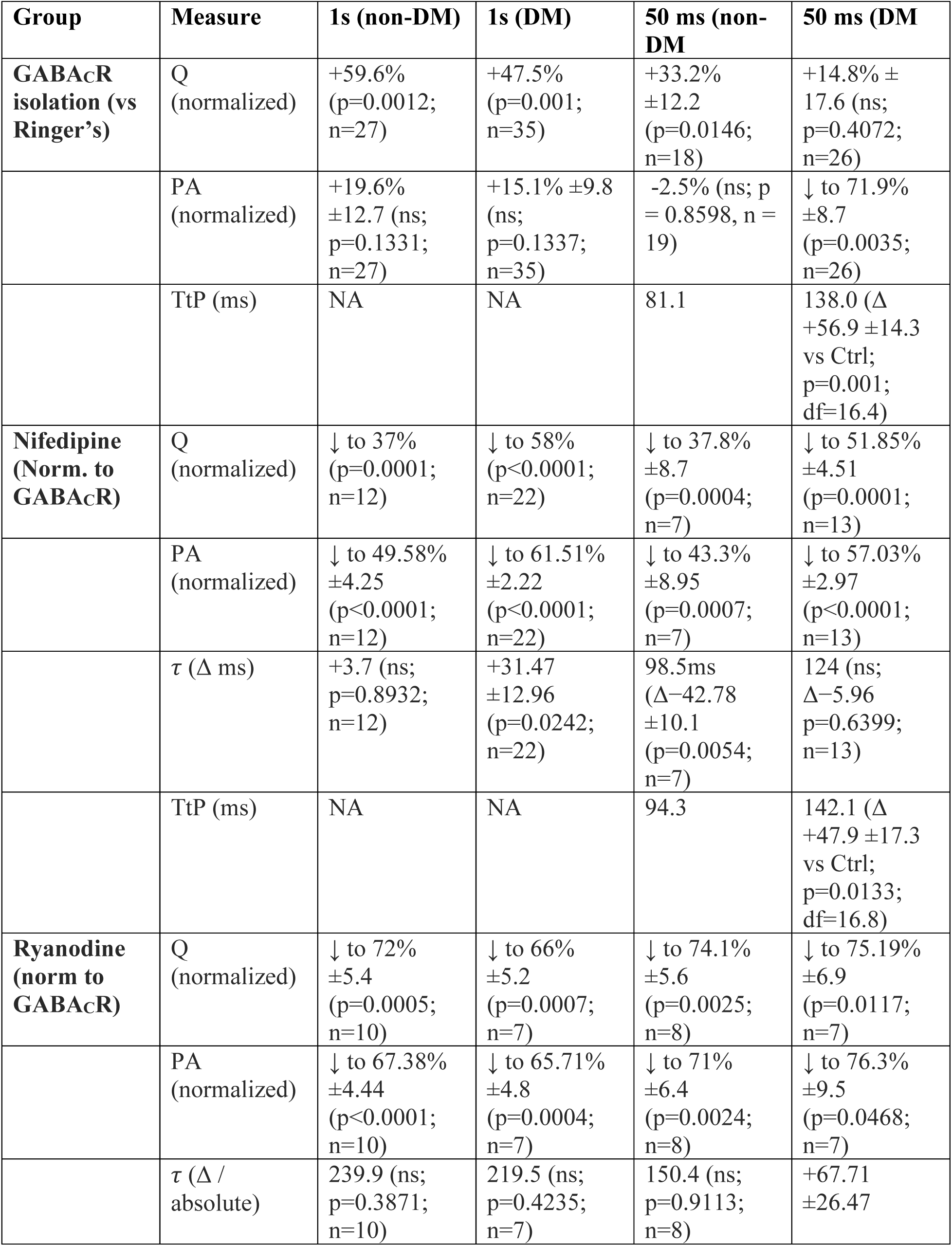
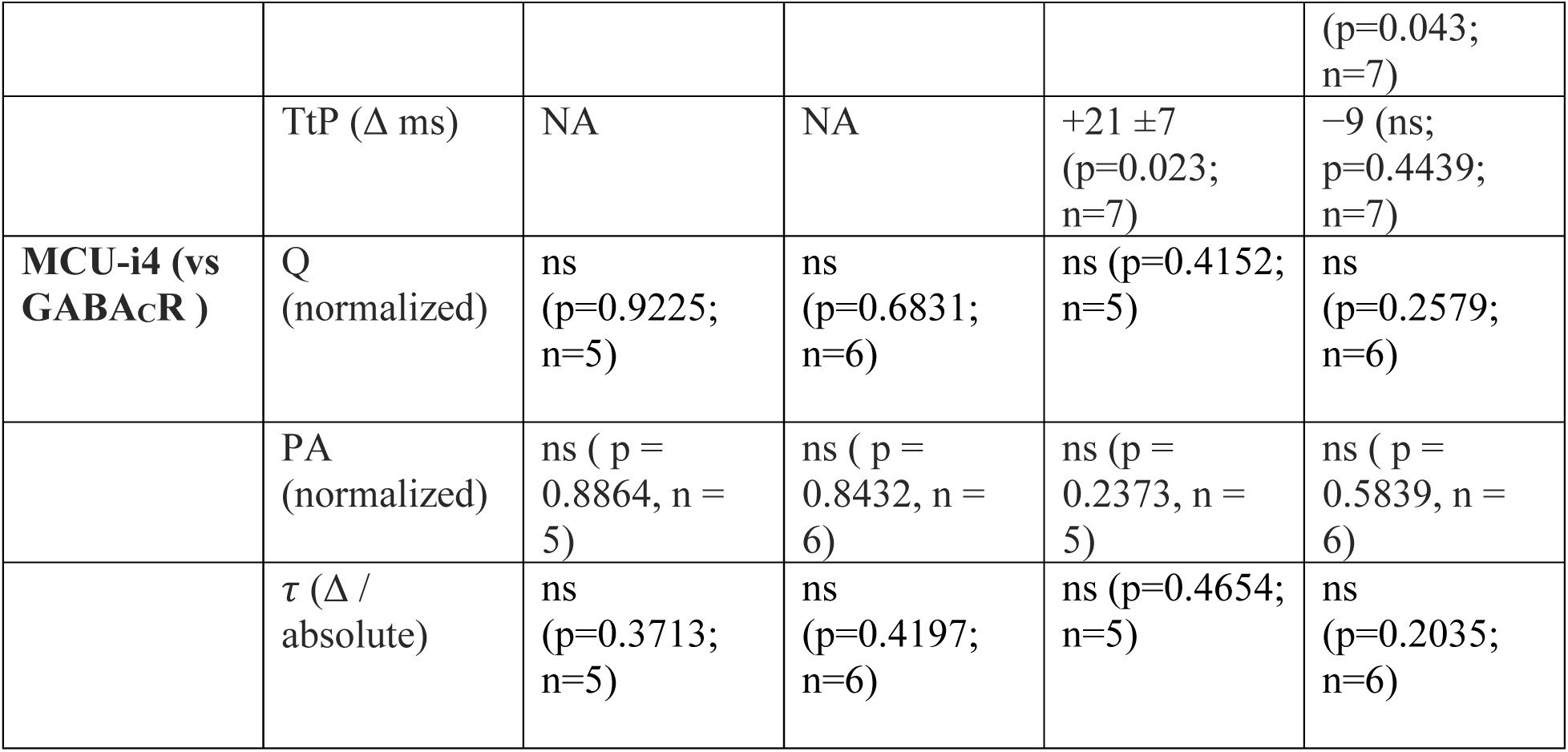

To test for the effects of the photoblock antagonists alone, we also measured the response to photoblock in a small data set in RBCs (Supplemental Figure 1). RBC responses are not significantly affected by the photoblock - owing to the fact that the pathway is fully light saturated. However, ON cone BC responses were significantly diminished by application of the photoblock, revealing that some of their response to the optogenetic blue light stimulus, included light activation of photoreceptor circuits (Supplemental Figure 1).

### Long optogenetic stimuli show changes in intensity and kinetics that are differentially affected by nifedipine in DM animals

The addition of nifedipine, an L-type Ca^2+^ channel blocker, significantly decreases these GABA_C_R isolated currents in non-DM and DM animals (Figure 2D). Between group comparison shows no significant difference in the mean strength of the nifedipine-diminished response (non-DM Mean Q = 12,275 pA*ms, DM Mean Q = 15,497 pA*ms, Difference between means = 3,222 +/- 5,313, p = 0.5483). The peak amplitude is significantly diminished by nifedipine in both groups (Figure 2C) but is significantly more diminished in the non-DM than in the DM group (Non-DM PA reduced to 49.58%, DM PA reduced to 61.51%, Difference between reductions = 11.93% +/- 4.62%, p = 0.0146). The more significant reduction of peak amplitude in the non-diabetic group suggests less tight coupling of release to the L-type Ca^2+^ channel in diabetes. This could be due to reduced CAV1.3 expression in DM presynaptic amacrine cells, which is suggested by IHC staining evidence (Figure 3A), showing decreased CAV1.3 overlap, with GAD65/67 in the DM inhibitory cells. Composite images with GAD, glycine, DAPI and CaV1.3 were taken with a confocal microscope and then multiplied against one another to find the overlap area (Figure 3C). The mean intensity of stains was no different between DM and non-DM (Figure 3B), suggesting no overall reduction in total CaV1.3 expression in the IPL. The significant reduction in overlap area between CaV1.3 and the inhibitory stains suggests that CaV1.3 expression may be primarily reduced in the inhibitory circuitry, but not throughout the rest of the IPL.

**Figure 2.**
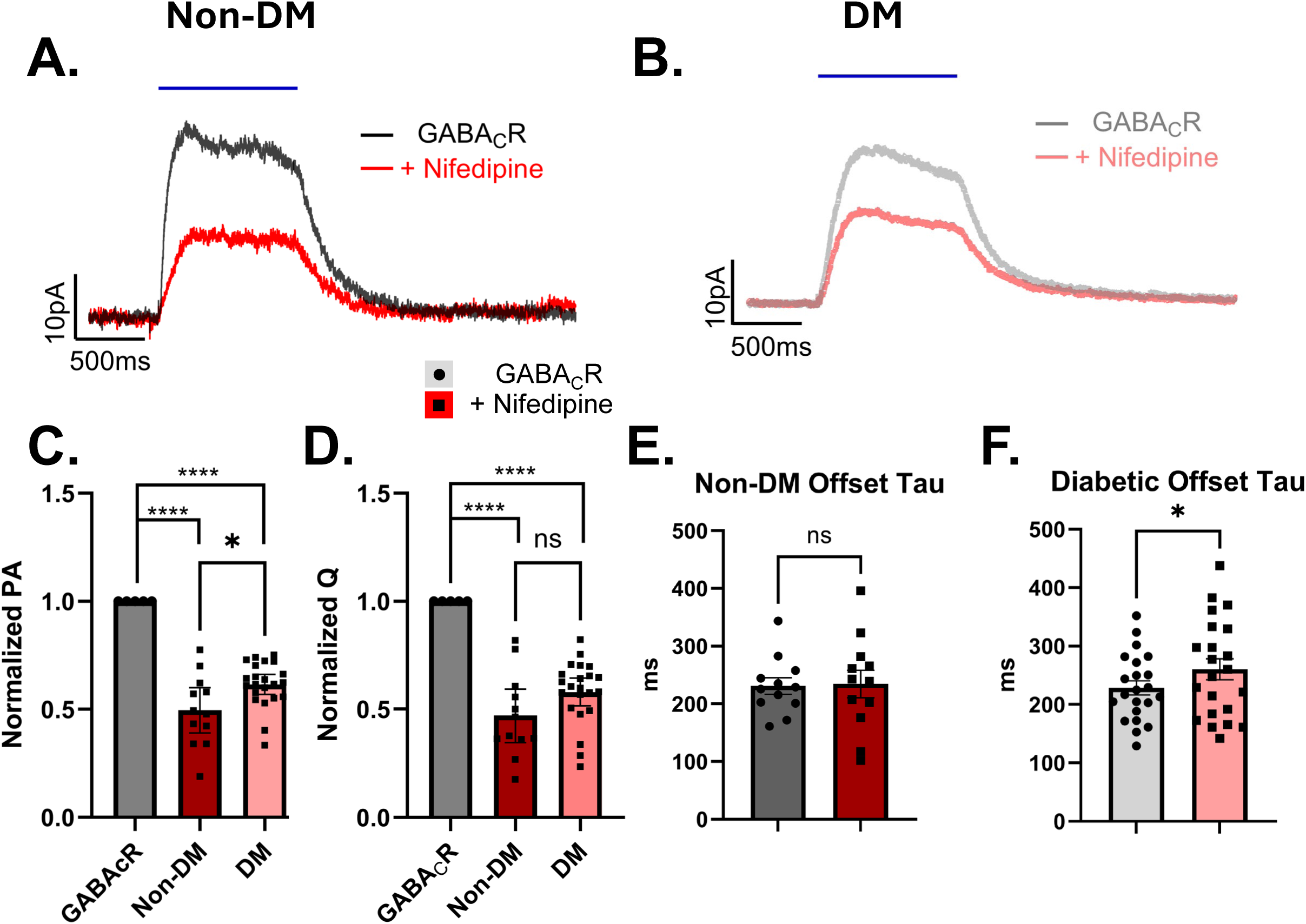
A, Averaged Non-DM traces from GABA_C_R represented by dark grey, application of Nifedipine represented by dark red. B, Averaged DM traces from GABA_C_R represented by light grey, application of Nifedipine represented by light red. Blue bar represents 1s optogenetic light stimulus for each. C, Peak Amplitude in the non-DM group is reduced more significantly (PA 49.58% +/-4.25%, p < 0.0001, n = 12) than it is reduced in the DM group (PA 61.51% +/- 2.22%, p < 0.0001, n = 22). There is a significant difference between groups (PA Difference = 11.93% +/- 4.6%, p = 0.0146). D, Both non-DM and DM responses diminish significantly for both non-DM (Q 47%, p = 0.0001, n = 12) and DM (Q reduced to 58%, p < 0.0001, n = 22) conditions. E-F, offset Decay Taus are not significantly different in the non-DM group (***τ*** +3.7 ms, p = 0.8932, n = 12), but are slightly increased in the DM group (***τ*** +31.47 +/- 12.96ms, p = 0.0242, n = 22). Peak amplitude and Q values are normalized to the GABA_C_R isolated value.

**Figure 3.**
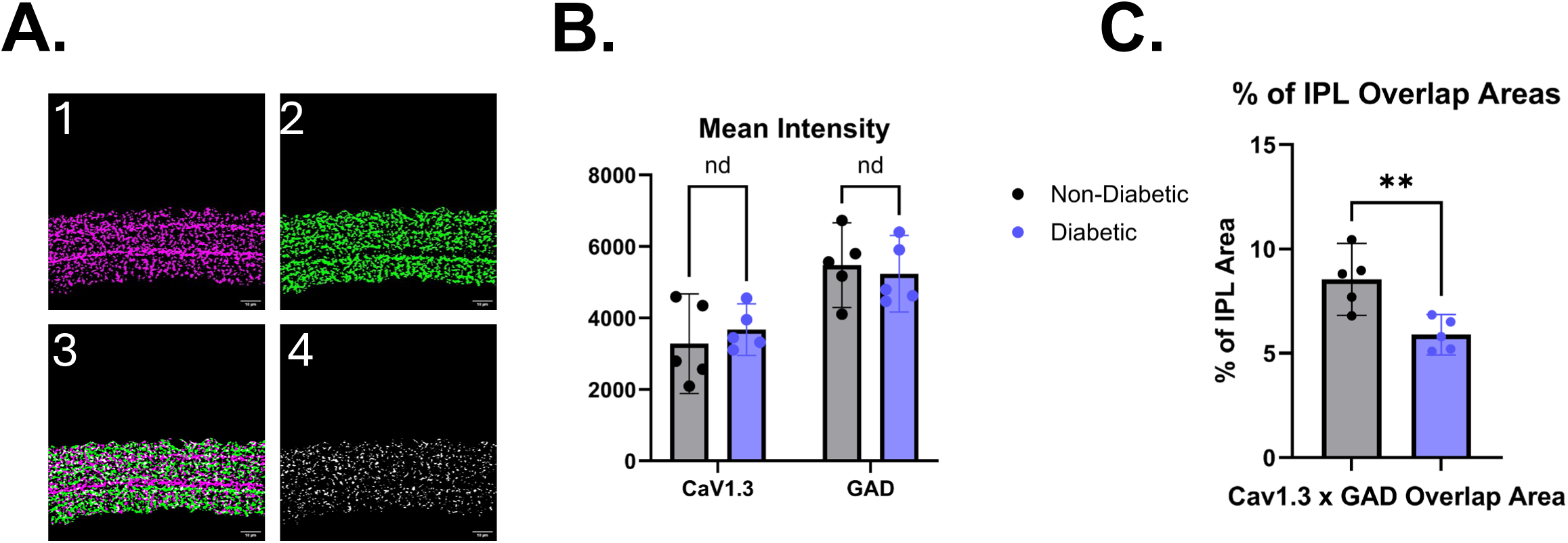
A, Image 1 depicts a thresholded z projection of the CaV1.3 stain (purple) in the IPL. Image 2, depicts GAD in green. Image 3 shows the overlaps of the thresholded z projections, with white representing overlap area. Image 4, Projections are then multiplied to distinguish overlap regions - pictured in white, lower right. B, Overall CaV1.3 and GAD stain intensity is not diminished in DM. C, in DM retinas Cav1.3 Channels have significantly less overlap with GAD - a marker for GABAergic cells (Difference between means = -2.65% +/- 0.7110%, df = 8, p = 0.0058).

Not only are intensity features of nifedipine-diminished GABA_C_R responses altered in early diabetes, but decay kinetics also diverge, with a small but significant increase in offset ***τ***observed in the DM group after nifedipine, which is not in the non-DM group (Figure 2E-F). This suggests that there is some degree of alteration in raw decay kinetics, but it may not be robust between groups. This result may represent an early DM change in intracellular Ca^2+^ handling, either in reduced clearance or buffering potential, which is revealed through blockade of L-type Ca^2+^ channels.

### GABA_C_R isolated peak amplitudes after brief stimuli are reduced in DM

In order to assess whether a shorter, more physiological stimulus condition could amplify circuit effects from diabetes, a 50ms stimulus was used. Traces taken as baseline recordings (only in the presence of Ringer’s solution) have very fast and sharp inhibition currents that are suppressed by GABA_C_R isolation. Similar to the 1s response, peak amplitude (Figure 4C, Table 1) is not significantly increased by GABA_C_R isolation in non-DM conditions, but charge transfer is significantly increased (Figure 4D). In the aggregated DM data, Peak Amplitude significantly decreases in the presence of GABA_C_R isolation (Normalized PA reduced to 71.93% +/- 8.72%, p = 0.0035, n = 26) but charge transfer is not affected (Figure 4C). Because the PA measure can include rapid glycinergic currents, this reduction in the DM condition could reflect DM alterations in either glycinergic or GABAergic inhibitory pathways.

**Figure 4.**
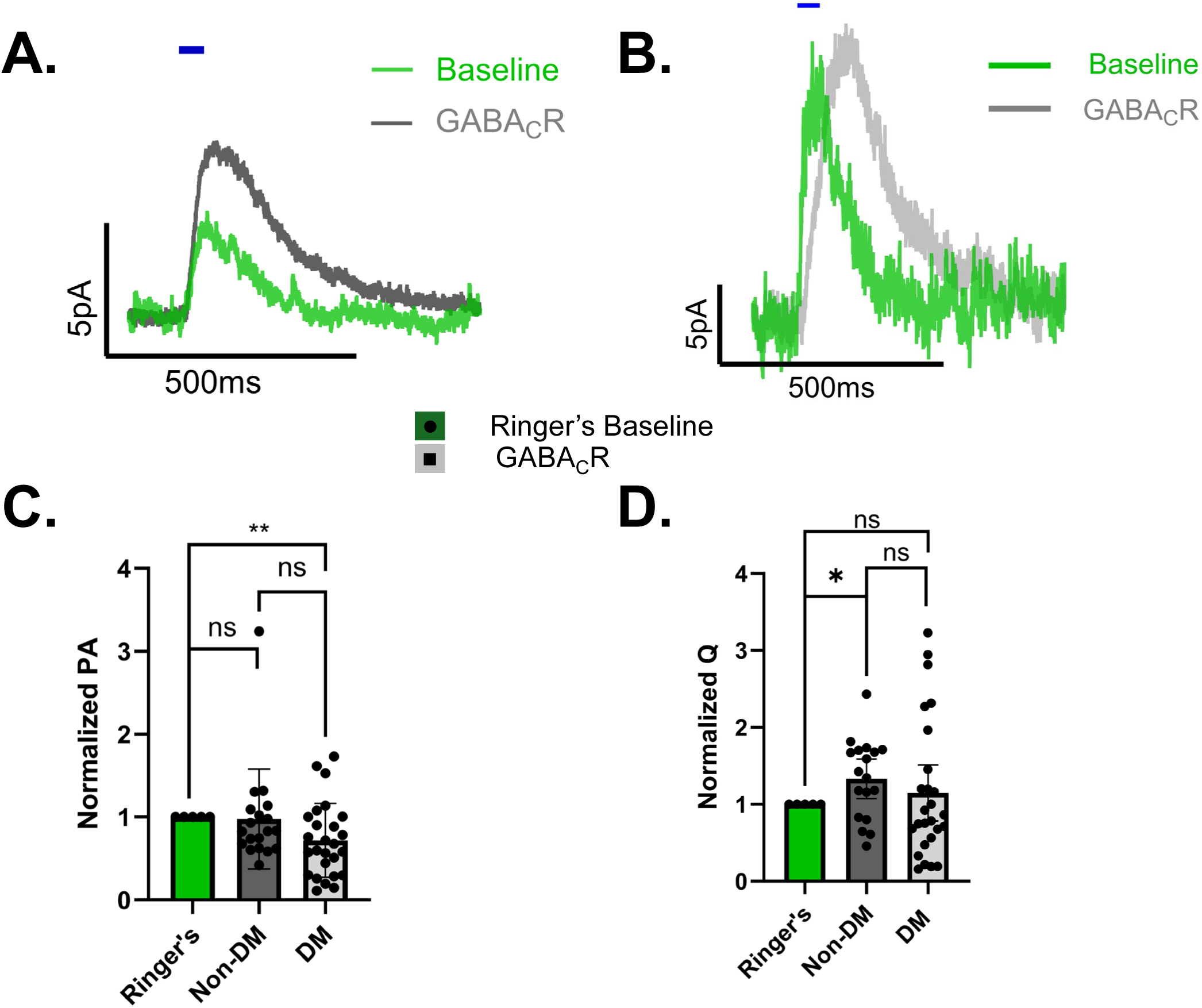
A, Averaged non-DM trace showing baseline Ringer’s recording in dark green, and GABA_C_R isolation in dark grey. B, Averaged DM trace showing baseline Ringer’s recording in light green, and GABA_C_R isolation in light grey. The blue bar represents the 1 second blue light stimulus for both. C, Non-DM peak amplitude is unchanged by GABA_C_R isolation (PA reduced to 97.5%, p = 0.8598, n = 19), while DM PA significantly decreases in the presence of GABA_c_R isolation (Normalized PA reduced to 71.9% +/- 8.7%, p = 0.0035, n = 26). Despite this, there is no statistically significant difference between DM and Non-DM groups (p = 0.1269, df = 31.6). D, Charge transfer significantly increases in Non-DM (Q +33.2% +/-12.2%, p = 0.0146, n = 18) but not for DM (Q +14.8% +/- 17.6%, p = 0.4072, n = 26), but there is not a significant difference between groups (p = 0.3958, df = 40).

### Nifedipine decreases decay timing from brief stimuli in non-DM, but not in DM animals

The response to nifedipine after a 50ms stimulus is very similar to the 1s stimulus data, where both PA (Figure 5D) and Q (Figure 5E) are significantly diminished by the presence of nifedipine, but without difference between experimental groups. Between group comparisons for Q are not significant (non-DM Q = 1949 pA*ms, Diabetic Q = 901.2 pA*ms, difference between means -1048 ± 655 pA*ms, p = 0.1530), and likewise with PA (p = 0.5011). The Decay Kinetics for the 50ms stimulus are altered in the non-DM group (Figure 4 C) with a significant decrease in ***τ*** (Nifedipine ***τ* =** 98.5ms, **Δ** from GABA_C_R -42.7ms ± 10.08, p = 0.0054, n = 7) but are unaffected in the DM group (Nifedipine ***τ* =** 124ms, **Δ** from GABA_C_R -5.96ms, p = 0.6399, n = 13) (Figure 5F).

**Figure 5.**
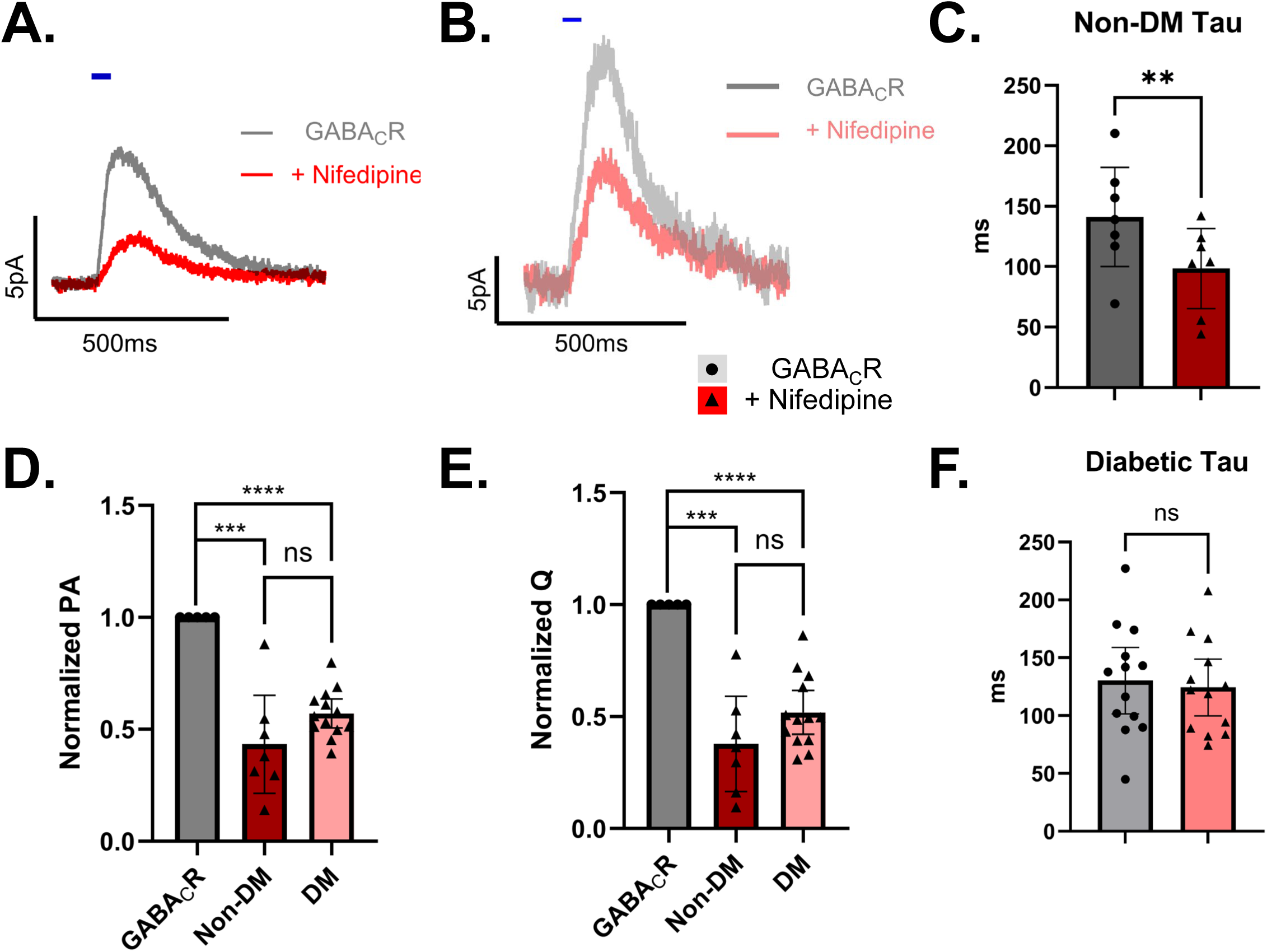
A, Averaged Non-DM traces from GABA_C_R represented by dark grey, application of Nifedipine represented by dark red. B, Averaged DM traces from GABA_C_R represented by light grey, application of Nifedipine represented by light red. The blue bar represents the 50ms second blue light stimulus for both. D, Peak Amplitude is significantly diminished in both groups. non-DM (PA ↓ to 43.3 +/ - 8.95% p = 0.0007, n=7) DM (PA ↓ to 57.03 +/- 2.97%, p < 0.0001, n = 13). E Charge Transfer is significantly and similarly diminished in both groups. (Q reduced to 37.8% +/ - 8.7% p = 0.0004, n=7) and in the DM (Q ↓ to 51.85 +/ - 4.51% p = 0.0001, n=13). C, in the non-DM group the decay tau is significantly lower(***τ* =** 98.5ms, Δ-42.78ms +/- 10.08ms, p =0.0054, n = 7), but it is unaffected in the DM condition**(*τ* =** 124ms, Δ-5.96, p = 0.6399, n=13).

With the shorter stimulus, Time to Peak (TtP) was a feature that we could meaningfully extract information from, and we have found that the time to peak is significantly increased in the DM group (TtP = 138ms) compared to the non-DM group (TtP = 81.1ms) for both GABA_C_R isolated traces and for Nifedipine traces (Table 1). Differences in both onset and offset kinetics in the 50ms condition suggest that the long 1s stimuli extracts separate features from the RBC-Amacrine circuit, and that the 50ms condition can provide relevant nuance to these altered Ca^2+^ handling dynamics.

### Ryanodine does not differentially affect non-DM and DM traces with long stimuli

To test the hypothesis that DM intracellular Ca^2+^ disruption occurs due to alteration in CICR, we used the potent ryanodine receptor antagonist Ryanodine at a concentration (50 µM) which locks the receptor into a closed conformation. This drug robustly decreases peak amplitude and charge transfer (Figure 6C-D) in each group. However, decay kinetics in 1s traces are not significantly altered by ryanodine (Figure 6E-F). This kinetics result is rather puzzling, as preventing ER stores from emptying should reduce the amount of calcium considerably and make it faster to return to baseline. In summary, 1 second DM traces show no statistically significant divergence from non-DMs, either in size or in kinetics, suggesting that CICR investigation through ryanodine is not relevant to the early disease progression of DM retinopathy when probed with the extremely long and powerful 1s optogenetic stimulus.

**Figure 6.**
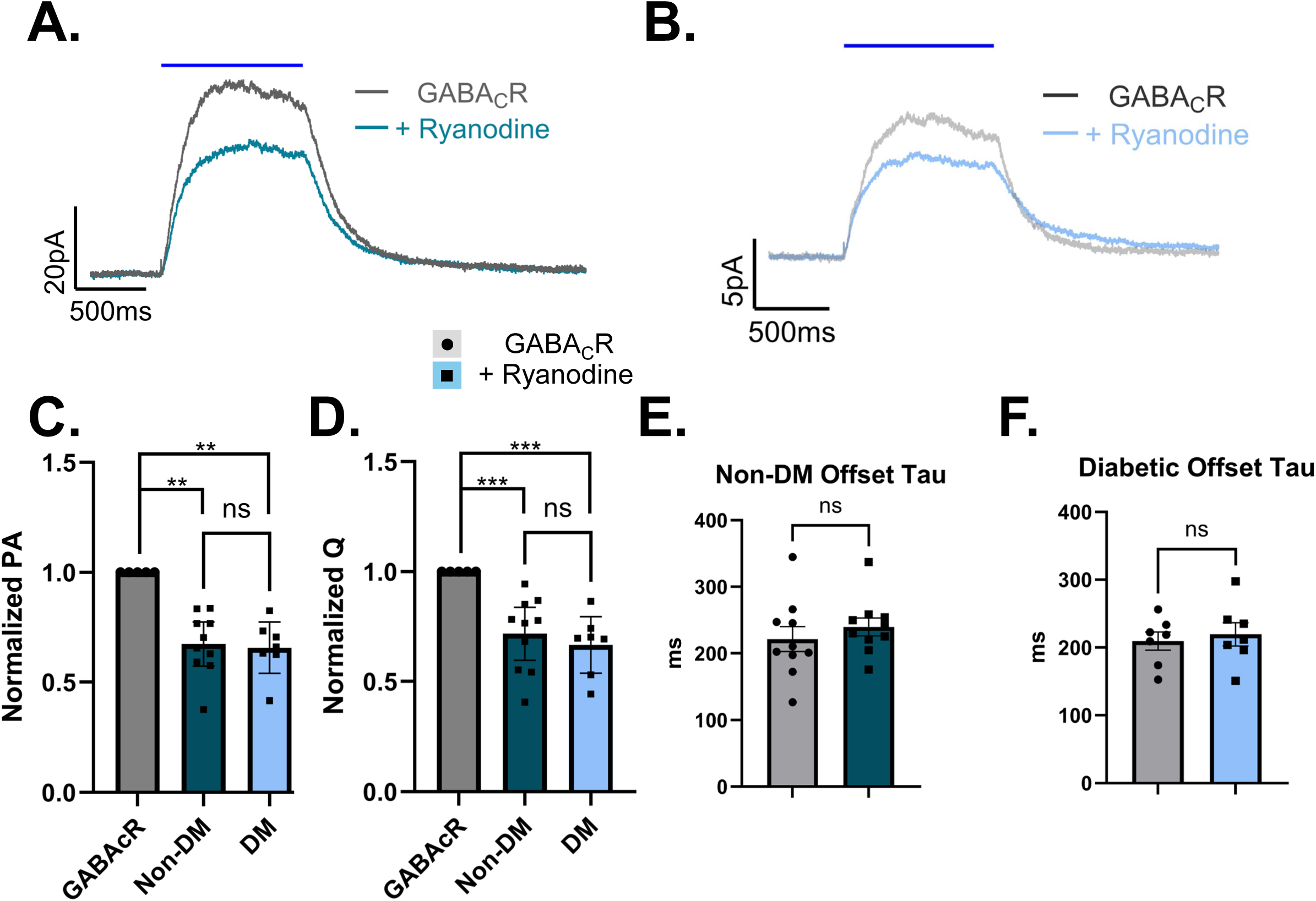
A, Averaged Non-DM GABA_C_R isolated traces represented by dark grey, application of Ryanodine represented by dark cyan. B, Averaged DM GABA_C_R isolated traces represented by light grey, application of Ryanodine represented by light cyan. Blue bar represents 1s optogenetic light stimulus for each. C, Peak Amplitude in the non-DM group is reduced (PA 67.38% +/-4.44%, p < 0.0001, n = 10) and is nearly equally reduced in the DM group (PA 65.71% +/- 4.8%, p = 0.0004, n = 7). D, Both non-DM (Q reduced to 72% +/-5.4%, p = 0.0005, n = 10) and DM (Q 66% +/-5.2%, p = 0.0007, n = 7) responses are diminished similarly by ryanodine. E,F the non-DM Offset Tau (***τ* =** 239.9ms, p=0.3871, n = 10) and DM Tau (***τ* =** 219.5ms p=0.4235, n = 7) are unaffected by ryanodine.

### DM affects the kinetics of ryanodine modulation of brief stimuli

Ca^2+^ induced Ca^2+^ release, which functions to amplify Ca^2+^ signals through the release of intracellular stores, could be thought to be most pronounced for a shorter stimulus. Theoretically, a long stimulus will deplete Ca^2+^ stores over time, leading to some dynamic equilibrium where the ER is relegated to the role of buffering out Ca^2+^, rather than amplifying the signal. Therefore, disruption of CICR at this much shorter time point should lead to a more pronounced reduction in the inhibitory response when compared to the 1s stimulus, as amplification will be shut off by the closure of ryanodine receptors. Responses were reduced to similar levels between groups in both normalized charge transfer and normalized peak amplitude (Figure 7C-D). These reductions are of similar magnitude as for the 1s stimulus, suggesting that CICR is equally relevant to the long and short stimulus.

**Figure 7.**
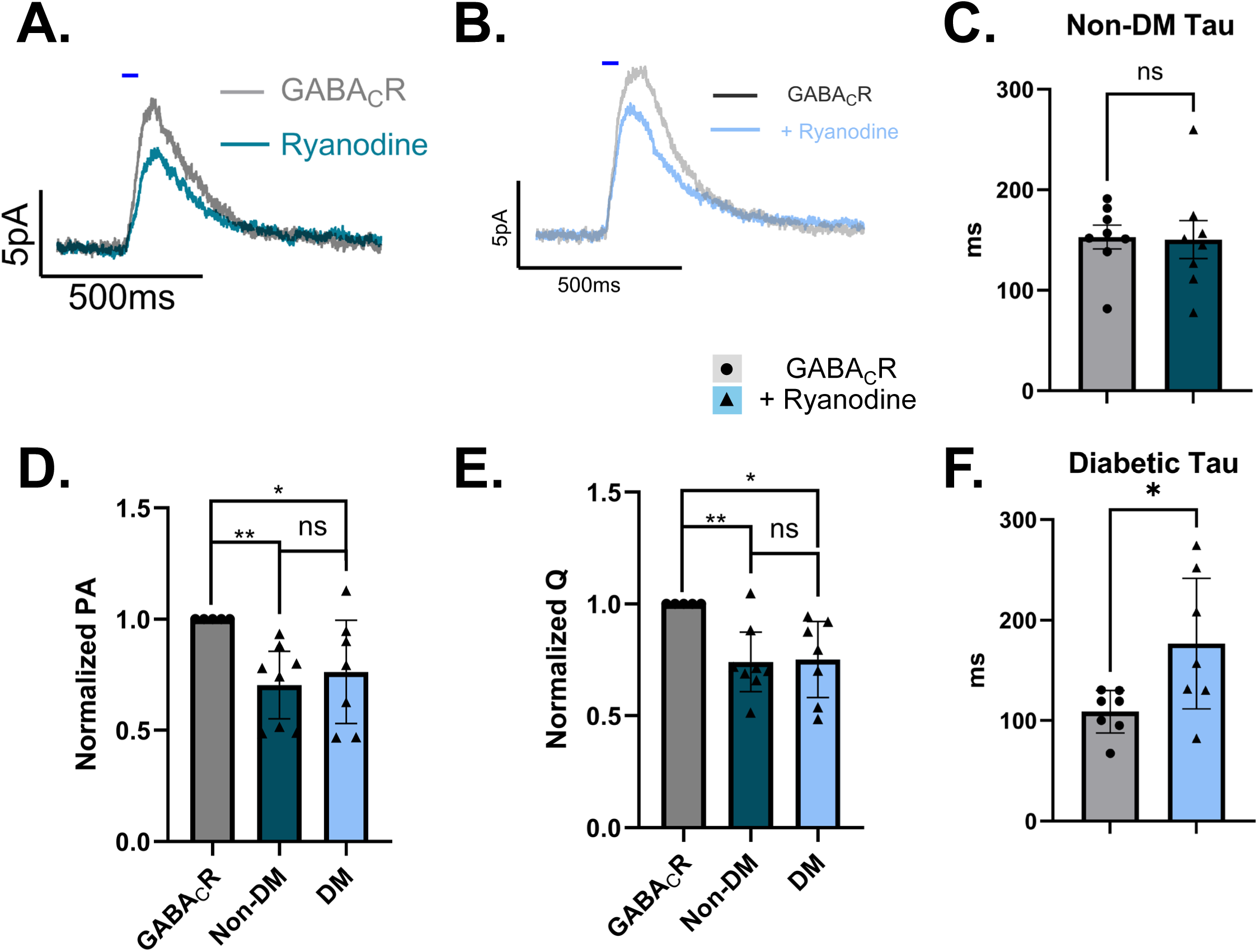
A, Averaged Non-DM GABA_C_R isolated traces represented by dark grey, application of Ryanodine represented by dark cyan. B, Averaged DM GABA_C_R isolated traces represented by light grey, application of Ryanodine represented by light cyan. Blue bar represents 50ms optogenetic light stimulus for each. D, PA is reduced for both non-DM (PA 71 % +/-6.4%, p = 0.0024, n = 8), and DM (PA 76.3 % +/- 9.5%, p = 0.0468, n = 7). E, The current is attenuated in non-DM (Q 74.1 +/- 5.6%, p = 0.0025, n = 8)) and DM groups (Q 75.19 +/- 6.9%, p = 0.0117, n = 7) by ryanodine. C, the non-DM Tau is unaffected by ryanodine (***τ* =** 150.4ms, p = 0.9113, n = 8). F, the DM tau is increased (***τ*** = 176.6, difference +67.71 +/- 26.47 ms, p=0.043, n = 7).

Unlike the 1s stimulus, kinetics change in the 50ms condition with a significant increase in diabetic decay tau in response to ryanodine (Figure 7 F), with no change in the non-diabetic group (Figure 7C). Based on the drug action of ryanodine this is not what we would expect but could reflect a source of dysregulation unveiled by CICR blockade. When comparing onset kinetics using time to peak data, there is a significant delay in TtP in the non-DM group (Table 1) but not in the DM group. Mechanistically, this is interesting because the blockade of ryanodine receptors should reduce the amplification of the response, making it more transient. Stochastic variability, or some form of compensation may explain this longer time to peak. Evidence for stochastic variability comes from the difference between TtP means between experimental cohorts, where the TtP for the DM 50ms stimulus was 138ms in the GABA_C_ nifedipine group, but is 101ms in this GABA_C_ condition, despite there being no difference in drug application. These groups should have the same mean TtP, but in this case must represent different populations or significant variance in the measure. Therefore, it is difficult to conclude that this delay in TtP represents a biologically active phenomenon and may be statistically anomalous.

In summary, while ryanodine application affects peak amplitude in both groups, decay and timing kinetics show high variability. These findings may reflect underlying inconsistency in Ca^2+^ handling among amacrine cells, separate population subtypes, or variability in circuit measurements, warranting future CICR specific studies.

### Mitochondrial Ca^2+^ Uniporter blockade does not significantly affect inhibitory signaling

The hypothesis that Ca^2+^ buffering by way of the MCU leads to alterations in Ca^2+^ homeostasis in diabetes was tested using the drug MCU-i4, which blocks the MCU. Ca^2+^ is shuttled from the cytosol of the cell into mitochondria where it is sequestered and contributes to the ionic gradient of the organelle. Therefore, blockade of this transporter should increase Ca^2+^ availability in the presynaptic terminal or reduce the Ca^2+^ buffering capacity of the cell, prolonging and increasing stimuli responses. In using this drug, however, no significant difference is observed in Q, PA, or Tau for either stimulus length (Figures 8–9). There does seem to be a bimodal distribution in the data, where some cells notability had a large increase in their response when in the presence of the MCU-i4. Other cells had responses that showed no change or were diminished in response to the MCU-i4, leading to an average effect that is not statistically different, in either the 1s or the 50ms stimuli conditions.

**Figure 8.**
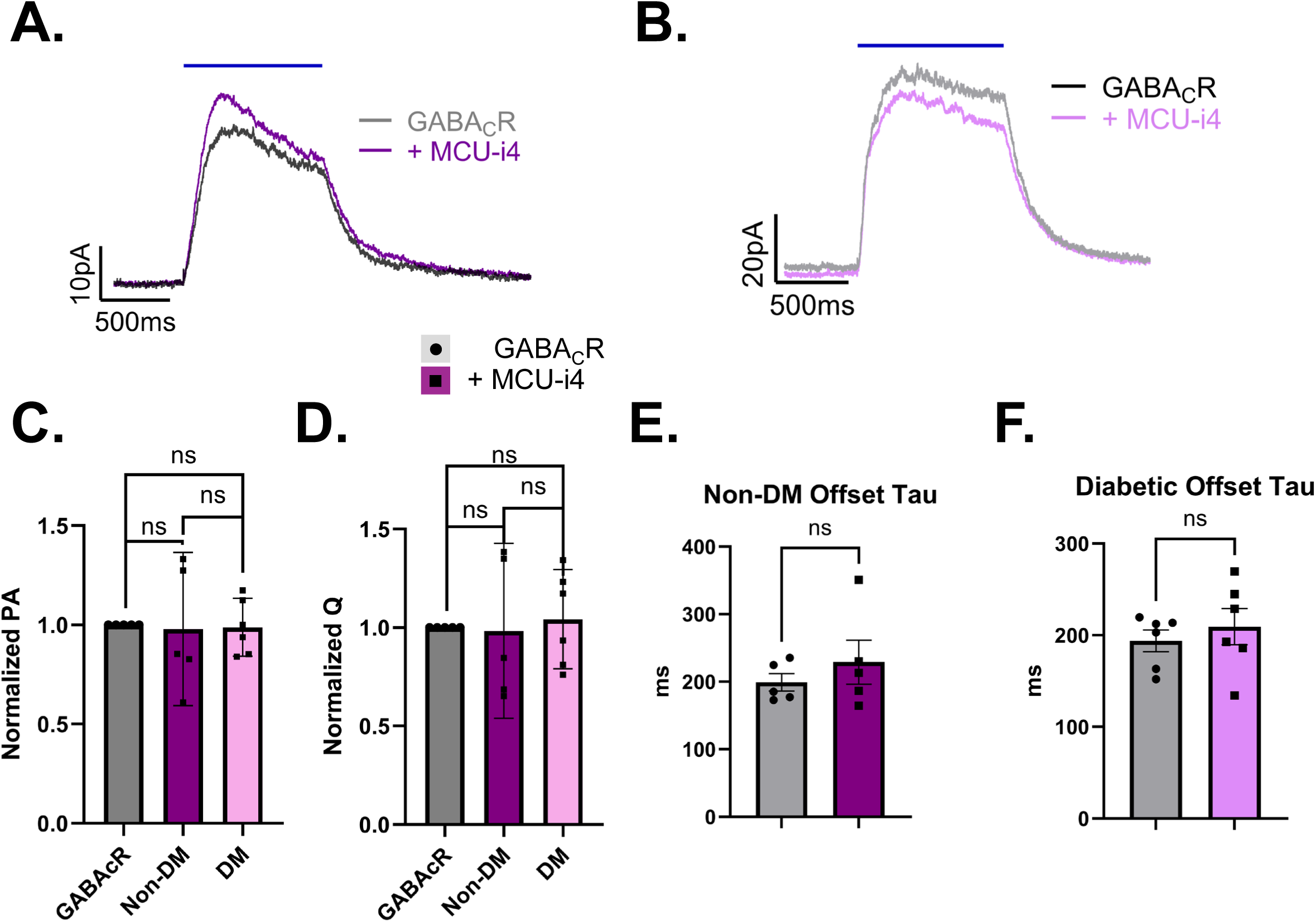
A, Averaged Non-DM GABA_C_R isolated traces represented by dark grey, application of MCU-i4 represented by dark purple. B, Averaged DM GABA_C_R isolated traces represented by light grey, application of MCU-i4 represented by light purple. Blue bar represents 1s optogenetic light stimulus for each. C, the peak amplitude is unchanged in Non-DM (PA p = 0.8864, n = 5) and DM (PA p = 0.8432, n = 6). D, The current is not attenuated in non-DM (Q p = 0.9225, n = 5) or in DM (Q p = 0.6831, n = 6). E-F, the non-DM and DM offset tau is unaffected by MCU-i4 (***τ***, p = 0.3713, n = 5), DM (***τ***, p=0.4197, n = 6).

**Figure 9.**
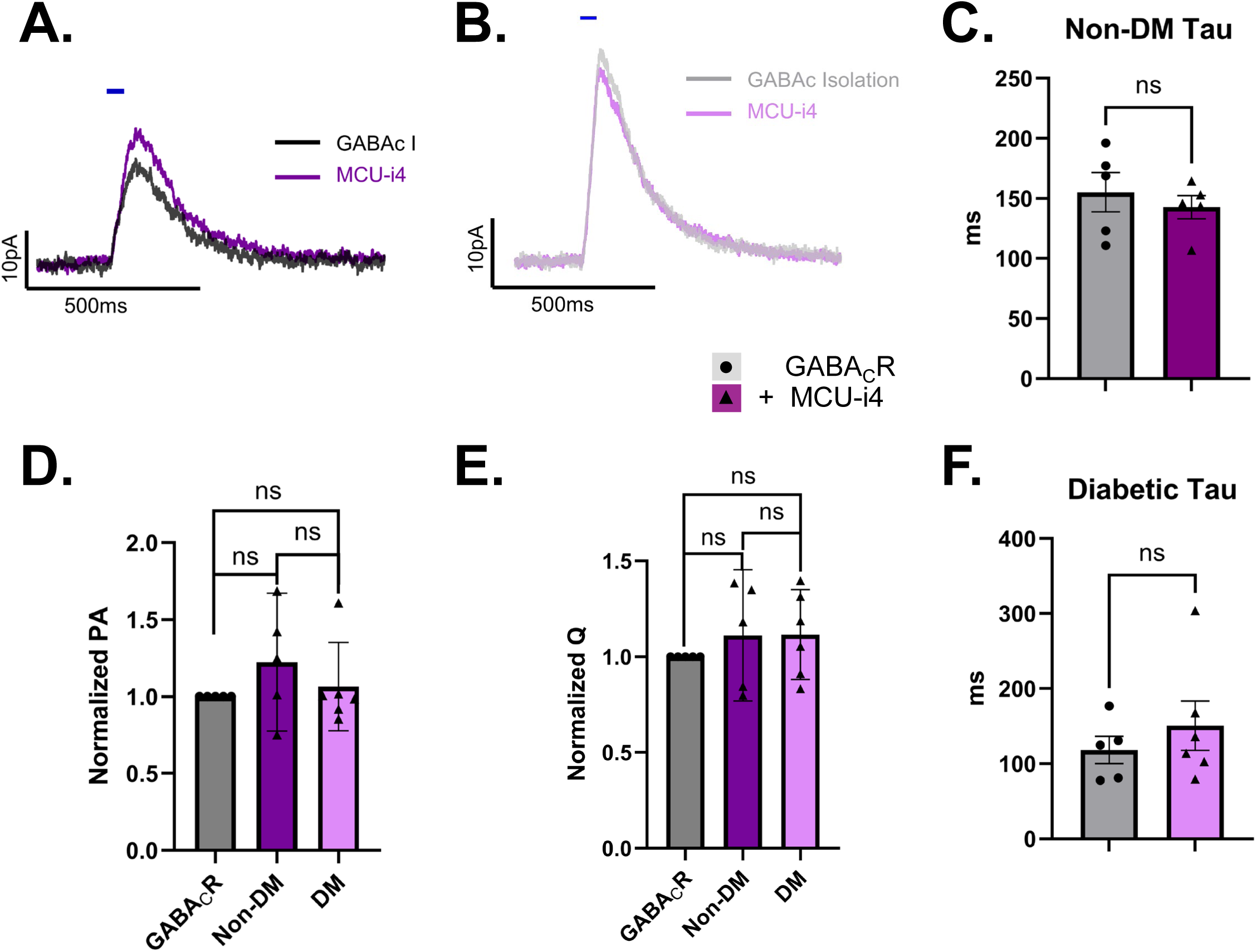
A, Averaged Non-DM GABA_C_R isolated traces represented by dark grey, application of MCU-i4 represented by dark purple. B, Averaged DM GABA_C_R isolated traces represented by light grey, application of MCU-i4 represented by light purple. Blue bar represents 50ms optogenetic light stimulus for each. D, peak amplitude is unaffected in Non-DM (p = 0.2373, n = 5) and in DM (p = 0.5839, n = 6). E, The current is not affected by drug condition in non-DM (Q, p = 0.4152, n = 5) or DM (Q, p = 0.2579, n = 6). C and F, the non-DM Tau is unaffected by MCU-i4 (***τ***, p = 0.4654, n = 5), as is the DM tau (***τ***, p=0.2035, n = 6).

These findings suggest that MCU-dependent buffering does not robustly regulate inhibitory output from amacrine cells in this circuit, or that only a subset rely on this buffering mechanism in these conditions. Future work is necessary to determine the source of this inconsistency, and the role of the MCU for synaptic regulation.

## Discussion

Diabetic retinal circuitry changes begin with alterations in amacrine cell output, as suggested by previous ERG and electrophysiology studies. This work serves to elucidate the mechanism for these changes by testing the hypothesis that early DM retinal circuitry is adjusting to a loss of Ca^2+^ tone in the amacrine cell, with mechanistic loci such as the voltage gated Ca^2+^ channel, ryanodine receptor, and the mitochondrial Ca^2+^ uniporter.

Overall, the recordings from these experiments provide evidence that L-type Ca^2+^ channel blockade unmasked Ca^2+^ dysregulation in diabetes, that CICR may be altered in its contribution to neurotransmission kinetics, and that mitochondrial Ca^2+^ uniporter blockade serves as a minor regulator of inhibition dynamics for oIPSC currents in RBCs.

These experiments present evidence that GABA_C_R isolated peak amplitude is decreased for DM 50ms oIPSCs (Figure 3), which may represent altered GABA_C_Rs or changes in presynaptic vesicle recruitment. Given that diabetes has known effects on PKC signaling, and that PKC can regulate GABA_C_Rs^19^, a postsynaptic effect cannot be ruled out here^20^. However, we previously showed that the peak of GABA_C_R spontaneous currents did not change in RBCs after 6 weeks of diabetes, so a change to GABA_C_Rs is unlikely^13^. This decrease in peak amplitude occurred relative to the Ringer’s baseline, which could also suggest that baseline response peak amplitudes were larger in diabetes, implicating the circuitry providing inhibition to GlycineRs or GABA_A_Rs as being overactive. Especially given that GABA_C_R isolation slightly increased charge transfer, it might be most likely that the DM baseline had particularly large, sharp transients that GABA_C_R isolation reduced.

The time to peak of these 50ms GABA_C_Rs oIPSCs was significantly elevated in the DM animal compared to non-DM (Table 1) which, in conjunction with the decrease in peak amplitude, suggests that inhibition via GABA_C_Rs is slower to onset in early diabetes. When nifedipine blockade is used, this trend is preserved, with relatively elongated decay taus (Table 1), and longer times to peak (Table 1).

Specifically, Nifedipine decreases the 50 ms stimulus Tau of RBC oIPSCs in the non-DM retina, but not in the DM retina. This decrease in the non-DM is consistent with previous studies, where blocking the L-type Ca^2+^ channels shortens responses to electrically evoked inhibition.

This makes theoretical sense, where a smaller influx of Ca^2+^ should be buffered out faster to reset to baseline. The lack of tau decrease in diabetes suggests that there is some source of Ca^2+^, or some response of the synaptic machinery to Ca^2+^, that is prolonged relative to non-DMs. This could represent the DM cell’s attempt to compensate for the loss of L-type Ca^2+^ influx. With the 1s stimulus, the apparent increase in decay tau suggests some source of dysfunction relating to Ca^2+^ buffering or sequestration. These results are not well described by a simple mechanism but can be rationalized through a multilayered series of observations.

It is well established by the Moore-Dotson paper, that DM cells are more sensitive to the Ca^2+^ buffer EGTA-AM, and therefore DM cells exhibit decreased Ca^2+^ levels, or increased Ca^2+^ buffering. In contrast, this study’s results suggest that DM cells are slower to buffer out Ca^2+^ after nifedipine is applied. Therefore, nifedipine is unmasking a feature of DM Ca^2+^ regulation. In the presence of high Ca^2+^ levels emergency buffering kicks on to prevent apoptotic signaling cascades, through MCU activity^21^. When the level of Ca^2+^ influx is decreased, however, the cell may rely more heavily on the capacity of normal buffering as through SERCA pump sequestration. If these SERCA pumps are sluggish^22^, or show reduced expression, then the tau of these responses would increase only after blocking a major component of the Ca^2+^ signal. This would be true of ryanodine blockade as well, where the reduction in CICR is enough to prevent the MCU from kicking online, and the slowness of regular buffering leads to an increase in tau for DM ryanodine cells (Figure 6). It is further possible that microdomains of Ca^2+^ concentration are altered in diabetes, explaining the more significant impact of EGTA-AM on DM cells. If there are weaker Ca^2+^ microdomains near to the synaptic machinery in DM cells, then a general buffer could preferentially reduce the DM signal, but less so for the non-DM, where healthy stores proximal to the machinery prolong release.

MCU activation pulls Ca^2+^ into the mitochondria in order to facilitate ATP production, and to temporarily balance cytosolic Ca^2+^ overload. The Ca^2+^ concentration in the mitochondria is generally in balance with the cell’s cytosol, and excess import is shuttled back out via the sodium Ca^2+^ exchanger in the mitochondria^21^. Defective Ca^2+^ uptake can result in enhanced Ca^2+^ due to loss of cytoplasmic buffering^23–25^.

Because DM cells exhibit impaired mitochondrial ionic gradients, proteins which regulate the MCU could be disrupted, or the NaXCa^2+^ may more quickly release Ca^2+^ back into the cytosol, potentially extending the duration of responses, and explaining the increase in tau observed in diabetes^21^. If mitochondrial buffering capacity is dysregulated in diabetes, then why did MCU-i4 not have a significant effect? Without paired sequential drug application, it may not be possible to unmask dysregulation when several features of the cell are potentially compensating for one another. If the MCU is contributing to these increased decay taus when nifedipine is applied, then using both the MCU-i4 drug and nifedipine in conjunction should clearly reveal the MCUs contribution.

## Conclusion

Determining a robust mechanism for DM Ca^2+^ dysregulation will require significant further study but promises to pull on a thread of cellular biology that could be highly relevant to cellular hyperglycemic adaptation broadly. Nifedipine has been shown to reduce inhibitory peak amplitude preferentially in non-DM retinas, and short stimulus response kinetics diverge in tau for the non-DM retina. DM TtP is prolonged regardless of nifedipine. These findings indicate altered amacrine cell calcium handling in early diabetes, revealed when L-type entry is blockaded. CICR contributes to response sizes at both durations, with significant but variable kinetic effects induced by ryanodine at 50ms. MCU-i4 blockade, though potentially mechanistically relevant, showed limited, or cell population specific contribution to calcium regulation. This work suggests that neurons are responding to the hyperglycemic insult of diabetes through complex alterations of their calcium handling machinery, and that further research into these changes is necessary for finding interventions into DM pathogenesis.

---

**Figure S1.**
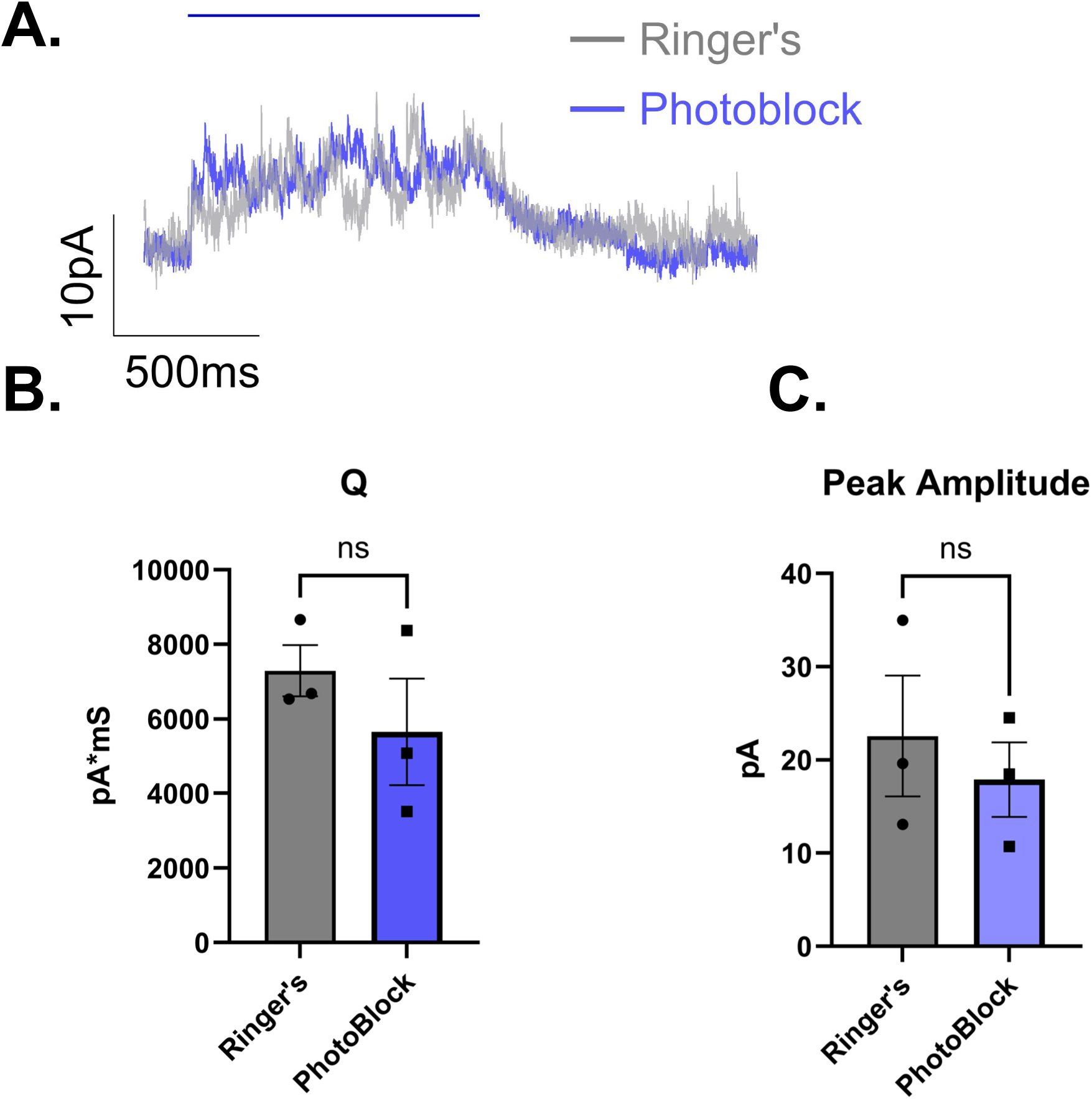
A, Sample trace from RBC, blue bar represents 1 second blue light stimulus. B - C, inhibitory currents were recorded from RBCs after ChR2 stimulation that were not significantly affected by the photoblocking cocktail (PA p = 0.34, Q p = 0.08, n=3).

**Figure S2.**
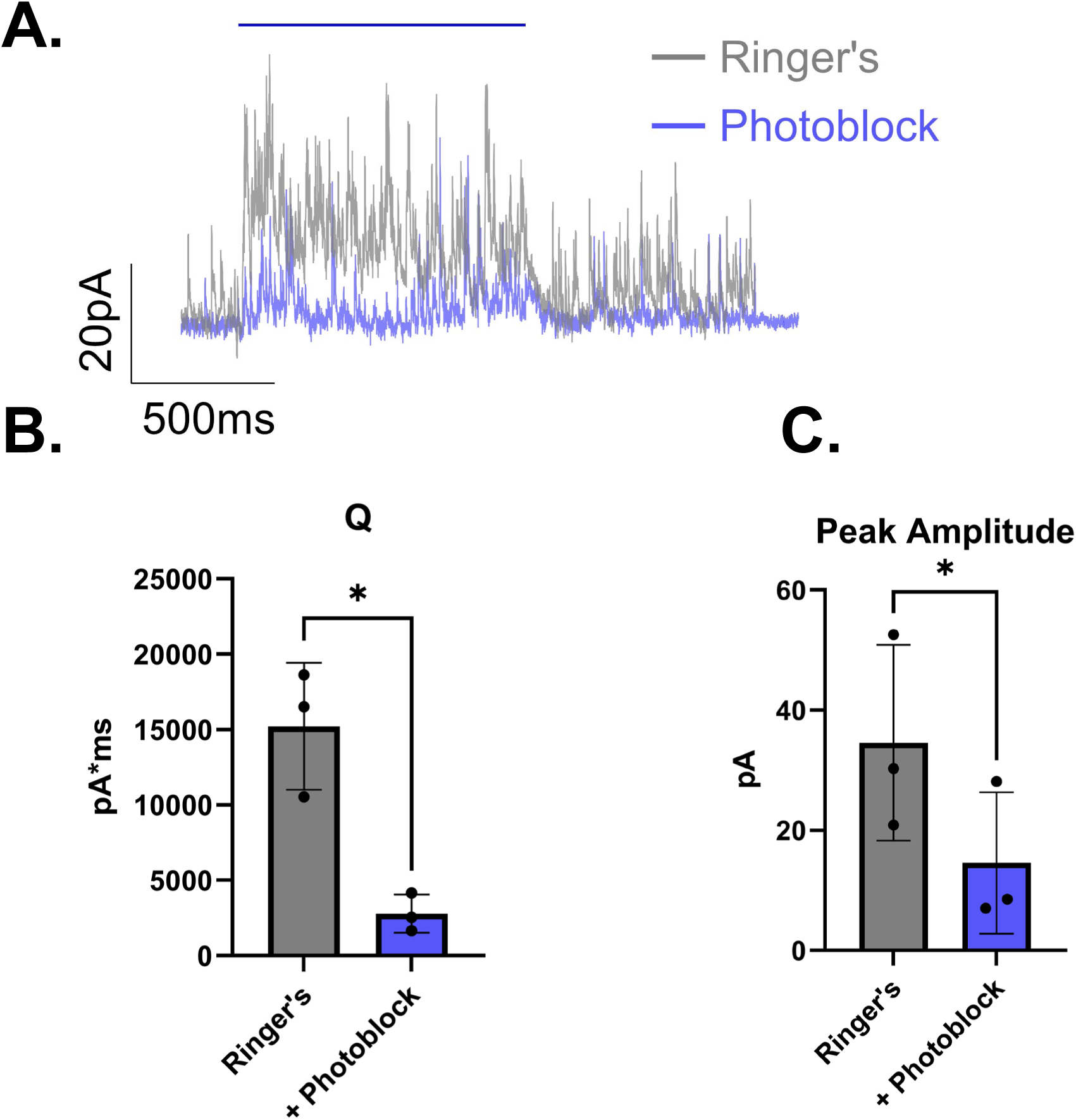
A, sample trace showing attenuation of the large current with application of the photoblocking cocktail. B-C the photoblocking cocktail significantly reduced ChR2 stimulated inhibitory currents in ONBCs (PA 58%, p = 0.025; Q 18%, p = 0.03, n = 3).

